# Standardized workflow for precise mid- and high-throughput proteomics of blood biofluids

**DOI:** 10.1101/2021.03.26.437268

**Authors:** Angela Mc Ardle, Aleksandra Binek, Annie Moradian, Blandine Chazarin Orgel, Alejandro Rivas, Kirstin E. Washington, Conor Phebus, Danica-Mae Manalo, James Go, Vidya Venkatraman, Casey Johnson, Qin Fu, Susan Cheng, Koen Raedschelders, Justyna-Fert Bober, Stephen R. Pennington, Christopher I. Murray, Jennifer E. Van Eyk

## Abstract

**Background:** Accurate discovery assay workflows are critical for identifying authentic circulating protein biomarkers in diverse blood matrices. Maximizing the commonalities in the proteomic workflows between different biofluids simplifies the approach and increases the likelihood for reproducibility. We developed a workflow that allows flexibility for high and mid-throughput analysis for three blood-based proteomes: naive plasma, plasma depleted of the 14 most abundant proteins, and dried blood.

**Methods:** Optimal conditions for sample preparation and DIA-MS analysis were established in plasma then automated and adapted for depleted plasma and whole blood. The MS workflow was modified to facilitate sensitive high-throughput or deep profile analysis with mid-throughput analysis. Analytical performance was evaluated from 5 complete workflows repeated over 3 days as well as a linearity analysis of a 5–6-point dilution curve.

**Result:** Using our high-throughput workflow, 74%, 93%, 87% of peptides displayed an inter-day CV<30% in plasma, depleted plasma and whole blood. While the mid-throughput workflow had 67%, 90%, 78% of peptides in plasma, depleted plasma and whole blood meeting the CV<30% standard. Lower limits of detection and quantitation were determined for proteins and peptides observed in each biofluid and workflow. Combining the analysis of both high-throughput plasma fractions exceeded the number of reliably identified proteins for individual biofluids in the mid-throughput workflows.

**Conclusion:** The workflow established here allowed for reliable detection of proteins covering a broad dynamic range. We envisage that implementation of this standard workflow on a large scale will facilitate the translation of candidate markers into clinical use.

## Introduction

Mass spectrometry (MS)-based proteomic analyses of blood provides a biochemical survey of an individual’s biological state.^1^ These insights can lead to an understanding of pathology and guide the development of clinically relevant biomarkers with diagnostic and prognostic utility for the outcome of a disease and the response to treatment.^1–3^ Blood and its constituent fractions (plasma and serum) are routinely used in clinical testing. Because of the minimal invasiveness of sampling, and the rich array of informative biomolecules they contain, serum and plasma are typically used in biomarker discovery studies. More recently, dried blood spots and volumetric absorptive microsampling (VAMS, i.e. Mitra^®^ microsampling device, Neoteryx, Torrance, CA) are emerging as alternative platforms for sample collection.^4,5^ Dried blood can be easily obtained by an individual at home without medical expertise or refrigeration, significantly expanding the potential for remote and longitudinal sampling, and for aiding analyses for difficult to access populations. While diversity in sample collection presents new opportunities for discovery and validation studies, it also underscores the need for an adaptable, scalable MS-based analysis platform that provides robust and reliable proteomic characterization, and that can facilitate the translation of new biomarkers to clinical use.

Common among the different blood biofluids is the dynamic complexity of their constituent proteomes. This complexity presents significant obstacles for robust and sensitive proteomic analyses. Liquid chromatography-mass spectrometry (LC-MS) can become overwhelmed by the highly abundant resident blood proteins and can fail to detect those less-abundant proteins that may signify a change in health status. Two archetypical strategies have been developed to overcome this challenge: (1) simplify the analyte by removing components that are unlikely to provide diagnostic value, and (2) optimize detection to make the survey of the biofluid more comprehensive. For example, simplifying blood to plasma or serum removes the cellular and platelet components; however, a small number of highly abundant proteins still dominate the analysis.^6^ These biofluids can be further simplified using selective depletion methods, for example the 14 most abundant proteins in plasma/serum,^7^ or targeted enrichment methods.^8–10^

To optimize the analysis of these sample types there are several key aspects to instrument set-up that can be considered. The choice of LC column, the flow rate, and the length of separation gradient can each drastically affect the quality of the analysis. Another key choice is the type of MS acquisition. In the past several years, data independent acquisition-MS (DIA-MS) approaches have gained preference to increase identification and quantitation of proteins in a complex sample. In DIA, all the precursor ions are fragmented across successive mass ranges to get a comprehensive survey of peptides in a sample. This can be achieved with MS instruments that have rapid duty cycle capabilities and high resolution. There are many MS instrument parameters like the size of the precursor mass windows that can be altered to improve protein depth and analytical precision in a DIA method.^11^

Although innovations in proteomic methodologies have emerged in recent years, there are still obstacles inhibiting the translation of biomarkers into clinical use. Chief among these are the requirement for advanced methods, highly trained personnel, and the costs associated with sample preparation and instrumentation. With these challenges in mind, we set out to establish a streamlined preanalytical proteomic workflow that achieves sensitive and robust analysis of multiple complex blood-based matrices, capable of quantifying the proteome of an individual across time, with sufficient reliability for large population-based screening.^12^ In this study, we present a robust standardized MS workflow that is intended to accommodate multiple blood biofluid inputs, sample processing, and throughput needs, in one unified analysis platform (Figure 1). In the following characterization, we present optimized digestion and acquisition conditions for naive and depleted plasma and dried blood and provide an inter- and intra-day assessment of the peptide and protein reproducibility and quantifiability performance. Using these proposed workflows, it is possible to balance throughput, proteome coverage, and cost to meet the needs of many biomarker discovery applications. Although we capitalized on using a state-of-the-art automation workstation (Beckman i7), our method can be easily executed in absence of any robotic workstation. The ability to switch between naive or depleted plasma and dried blood samples using the scalable workflow while achieving analytical precision is a necessity to develop translatable clinical biomarkers.

**Figure 1.**
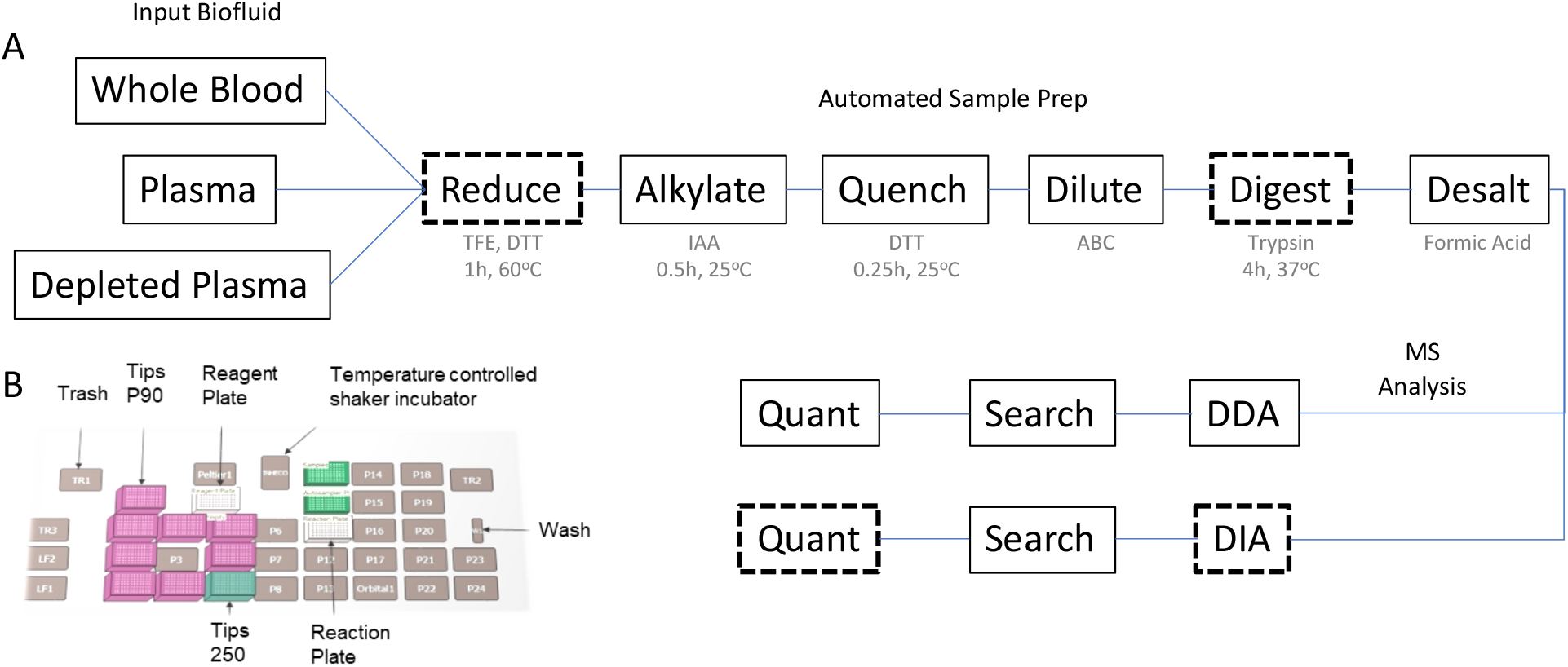
Standardized Workflow for Proteomic Analysis of blood biofluids. (**A**) Overview of steps required for preparation and analysis of plasma and blood. Dashed outline indicates a point of optimization in the procedure. (**B**) Schematic representation of automation platform.

## Methods

### Test Blood Biofluid Samples

Commercially available pooled mixed sex plasma (K2EDTA) was used for both analysis of naive and depleted plasma. Whole blood (K2EDTA) was purchased from Bioreclamation (Hicksville, NY, US) and mock collected using a Mitra^®^ microsampling devices (Neoteryx, Torrance, CA, US). See online supplement for more details.

### 96-well Format Top 14 Abundant Protein Depletion for Plasma

Plasma samples were depleted of its 14 most abundant proteins, albumin, Immunoglobulins A, E G and M (kappa and lambda light chains), alpha-1-acidglycoprotein, alpha-1-antitrypsin, alpha-2-macroglobulin, apolipoprotein A1, fibrinogen, haptoglobin, and transferrin using the High Select Top 14 Abundant Protein Depletion Camel Antibody Resin (Thermo Fisher Scientific). On the day of depletion, anti-camel antibody-resin, which was stored at 4 °C, was equilibrated to room temperature for 30 min mixing at 800 rpm. After equilibration, the anti-camel antibody-resin was vortexed vigorously and 300 ul was aliquoted into the wells of a 96 well plate (Nunc™ 96-Well Polypropylene DeepWell™ Storage Plates). 10 ul of plasma was diluted 1:10 with 100 mM NH_4_CO_3_ and added to wells containing depletion resin. To ensure homogenous mixing the plate was mixed at 800 rpm for 1 hour (hr). The unbound fraction was aspirated from the resin with 500 ul of 100 mM NH_4_CO_3_ and transferred to a filter plate (Nunc™ 96-Well Filter Plates). The depleted fraction was collected by gentle centrifugation (100 rcf for 2 min) into a clean 96 well plate (Beckman Coulter, deep well titer plate polypropylene) and lyophilized.

### Automated Blood Biofluid Trypsin Digestion and Desalting

All blood biofluids underwent protein denaturation, reduction, alkylation, digestion using the Beckman i7 automated workstation (BeckmanCoulter) programed for uniform mixing as previously described at a controlled temperature and modified with online automated desalting.^13^ Proteins were denatured in a solution of 35% v/v 2-2-2 Trifluoroethanol (TFE, Sigma), 40 mM Dithiothreitol (DTT, Sigma) and dissolved in 50 mM NH_4_CO_3_ (Sigma). The sample was denatured for 1 hr at 60 °C. Samples were then alkylated for 30 min at 25 °C in the dark with the addition of iodoacetamide (Sigma, (10 mM final concentration). To prevent over-alkylation DTT was added at final concentration of 5mM and samples were incubated for a further 15 min at 25 °C. Next, a volume of 50 mM NH_4_CO_3_ was added to dilute out TFE to a final conc of 5 or 10%. Trypsin was added to a ratio of 25:1 and samples were incubated for 4 or 16 hr at 37 or 42 °C. Digestion reactions were quenched with 5 ul of 25% FA. Desalting was carried out using a positive pressure apparatus (Amplius Positive Pressure ALP, Beckman Coulter) mounted on the left side of the i7 workstation deck. See SuppFig1 and the online supplement for more details.

### High- and Mid-throughput DIA LC-MS/MS

DIA analysis was performed on an Orbitrap Exploris 480 (ThermoFisher, Bremen, Germany) instrument. Our high-throughput workflow utilized the Evosep One system (Odense, Denmark) with a 21-min gradient requiring 25 mins to complete each sample. The mid-throughput workflow was run on an Ultimate 3000 ultra high-pressure chromatography system with a 45-min. gradient requiring 60 mins to complete. See online supplement.

### Bioinformatic Data Analysis

A full description of the data analysis and all peptide and proteins identifications including reproducibility and linearity characterizations can be found in the supporting information and Supplemental Tables 1-14. The coefficients of variation (CV) for protein or peptide intensities were determined if there were at least 3 out of 5 observations on each day. This threshold was required for all 3 days to determine a multi-day CV. Observed lower limit of detection (LLOD) for a protein or peptide based on the linearity experiment and determined by the lowest concentration of sample where the protein or peptide is detected in all 3 replicates with a CV<20%. Observed lower limit of quantitation (LLOQ) for a protein or peptide, was determined by the lowest concentration of sample where the protein or peptide is detected in all three replicates with an r^2^>0.8, CV<20% and a target deviation>0.2.

## Results

### Optimization of Digestion Parameters for Efficient Proteolysis in plasma

We assessed the impact of temperature and time on trypsin proteolytic efficiency (Figure 2A) using manual sample preparation. Plasma was digested with trypsin at 37 or 42 °C and analyzed by MS (n=3). Pearson correlation demonstrated that the relative peptide intensities from DDA analysis of all replicates were highly reproducible regardless of incubation temperature used; 37 °C (avg r^2^ =1.00) *vs.* 42 °C (avg r^2^ =0.99) (Figure 2B). To reduce sample processing times, we evaluated the impact of trypsin incubation length. Plasma was digested for 4 or 16 hr and analyzed by MS (n=3). The data revealed that using a 16 hr incubation period yielded a slightly greater (0.8-fold) peptide intensity response versus 4 hr (Figure 2C). Additionally, the longer incubation (16 hr) was associated with greater (0.8-fold) peptide identifications (average spectral counts 12,389) in comparison to the shorter time (average spectral counts 10,401). However, even though the 16 hr digestion displayed more favorable data in terms of number of identifications and relative intensity values, the duration of proteolysis did not impact on reproducibility. Indeed, relative intensity values across replicates could be highly correlated with average r^2^ value of 1.00.

**Figure 2.**
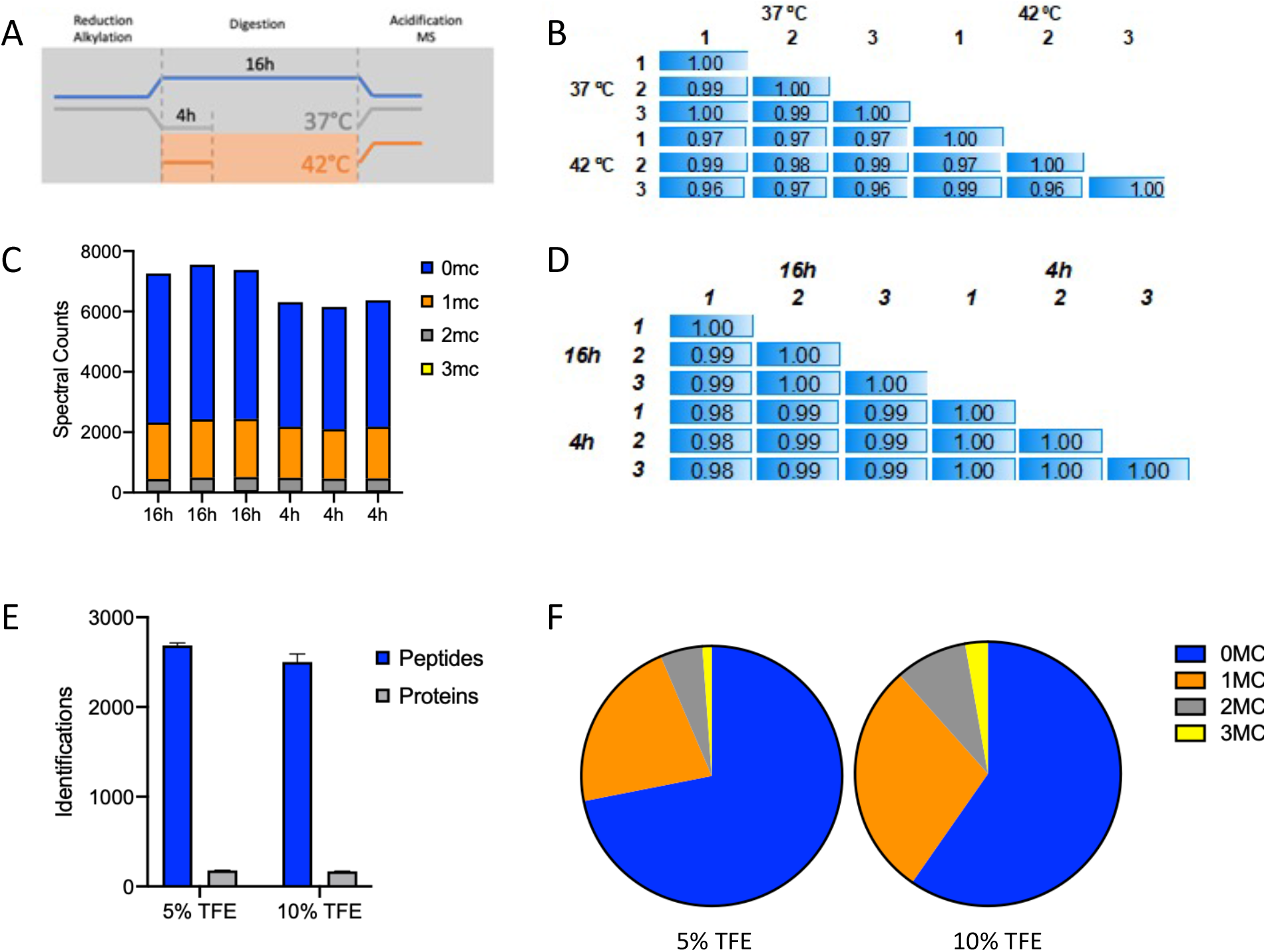
Optimization of digestion parameters. Digestion parameters optimization scheme (**A**) Peptide quantitation correlation for different settings in plasma digestion time (4 vs. 16 hr, n=3) and incubation temperature (37 °C vs. 42 °C, n=3) in blood and plasma. While correlation in plasma peptides quantitation was similar in both incubation times, it was improved at 37°C with respect to 42 °C (**B**), Log2 transformation of MS1 peak areas of peptides with 0, 1, 2 or 3 miscleavages (MC) in each digestion replicate at 4 and 16hr of incubation (**C**), Spectral counts of peptides with 0, 1, 2 or 3 MC at 4h and 16h of incubation. Recovery from digestion is proportionally similar in both incubation times. (**D**), Yield of peptide and protein identification under different denaturation buffers conditions (TFE 5% vs. 10%) (**E**), Number of peptides with 0, 1, 2 or 3 MC recovered from digestion when using TFE 5% or TFE 10% denaturation buffers. Blue, red, grey and yellow show proportion of peptides with 0, 1, 2, and 3 MC. Improved recovery from digestion was observed under denaturation conditions with TFE 5% (**F**).

The volatile denaturant, TFE, was used to improve digestion efficacy; however, the final concentration can influence the pH of the digestion buffer and impact trypsin activity. Analysis of plasma digested in 5% or 10% (v/v) TFE revealed that 5% TFE resulted in a marginal increased number of peptide identifications (2,686) compared to 10% (2,502). This observation can be attributed to the decreased rate of peptide miscleavages, a surrogate measure for proteolytic efficiency.^14^ In proteolysis with 5% or 10% TFE, 72% and 59% of peptides were fully cleaved, respectively. Reducing miscleavages is key as the signal loss from inconsistent digestion can lead to dubious relative intensity values (Figure 2F). To maximize throughput and efficiency, we selected conditions; 4 hrs. trypsinization at 42 °C in the presence of 5% TFE for inclusion into our final protocol.

### Optimization of DIA Acquisition Parameters for Plasma

Based on previous reports in plasma^15^ we compared the performance of 8 MS methods covering 4 isolation window widths and 2 resolutions (15 Da × 77 windows, 18 Da × 54 windows, 21 Da × 50 windows, 40 Da × 20 windows, each at 15,000 or 30,000 MS2 resolution). The number of protein identifications across all parameters were comparable, ranging from an average of 173-206 identifications. The data demonstrated that the 15 Da × 77 isolation windows at 30,000 resolution method supported the most peptide identifications compared to the other settings assessed (Figure 3B and 3C). Despite being associated with the greatest number of peptide identifications this method performed worst in terms of reproducibility (144 proteins, CV<30%, n=3). We found that the 15 Da method using 15,000 resolution supported fewer peptides (569) quantifiable with a CV<30% compared to the other methods which ranged between 654-841 peptides. Importantly, the 21 Da method using 15,000 resolution was associated with the greatest number of quantifiable peptides (841) with CV<30%.

**Figure 3.**
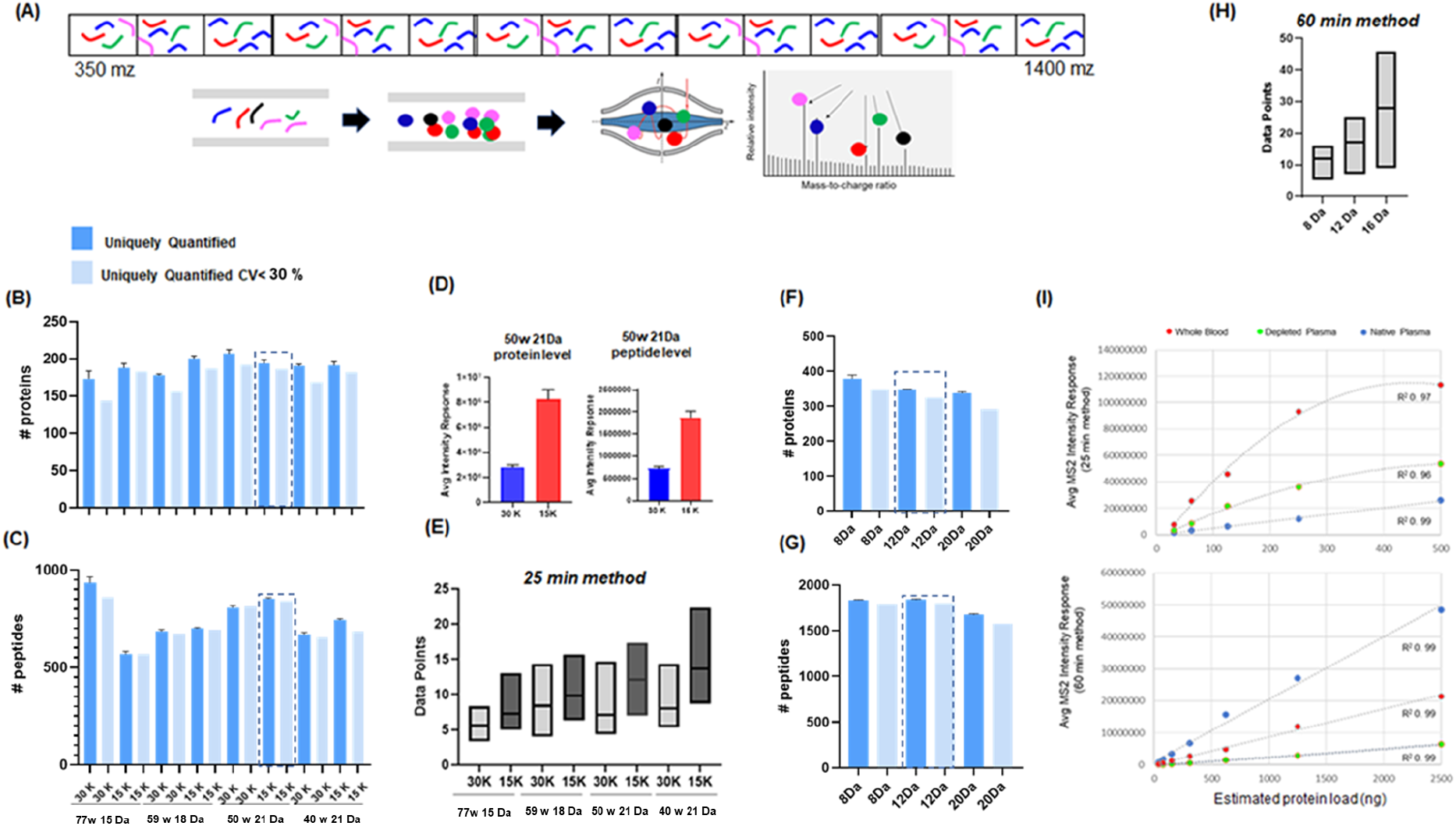
Optimization of DIA – MS for rapid, robust, deep and sensitive proteomic analysis of plasma. (**A**) In DIA-MS peptides that fall within a defined mass window are fragmented and filtered through to the orbitrap where fragment ions are analyzed simultaneously. The instrument cycles through subsequent overlapping windows until the entire mass range is covered. The instrument cycle time is influenced by resolution settings and number of mass windows which in turn impact on the number of identifications and reproducibility. (**B-C**) The number of proteins and peptides uniquely identified and quantified with CV<30% when analyzed with 8 different MS settings and a 25-min method (1 μl/min) (**D**) The averaged intensity response of all proteins and peptides when analyzed with resolution setting of 15,000 vs 30,000 (21 Da method). (**E**)The average number of data points under the curve from 11 iRT standard peptides when analyzed using 25-min method and 8 different MS2 settings. (**F-G**) The number of proteins and peptides uniquely identified and quantified with CV<30% when analyzed with 60-min method (9.5 ul/min). (**H**) The average number of data points under the curve from 11 iRT standard peptides when analyzed using 60-min method and 3 different MS2 settings.

Next, we investigated the impact of MS resolution settings, 15,000 or 30,000, on overall plasma protein and peptide intensity. We calculated the average intensity response from all identified proteins and peptides analyzed with 21 Da method employing either 15,000 or 30,000 resolution and found that 15,000 yielded the greater intensity response (Figure 3D). We hypothesize that this observation was attributed to the number of data points acquired across the curve for all detected peptides. Focusing on the 11 iRT peptides spiked into the plasma matrix, we plotted the minimum, maximum, and average data points across the curve observed when analyzed using the 8 MS parameters described above. As anticipated, the data revealed that 15,000 resolution supported more data points compared to 30,000 resolution (Figure 3E). The 40 Da windows method promoted the most data points (14.8±4.8, n=3) while the 15 Da at 30,000 resolution method supported the least data points (5.0±2.6, n=3). The 21 Da at 15,000 method performed second best in terms of the number data points collected (13.0±2.7, n=3). Importantly, data acquired using this method were associated with >8 data points across the curve, a well-established benchmark for acceptable quantification.^16^

Leveraging the results from the high-throughput workflow, we compared 3 DIA methods for our 60-min mid-throughput, workflow. Since this platform supported better peptide separation, we designed DIA methods with narrower isolation window settings and investigated the use of 8, 12 and 20 Da isolation windows. Here the broadest window setup (20 Da) was associated with the lowest number of proteins (292) and peptides (1,576) with CV<30%. While the data indicated the 12 Da method was the best performer (325 proteins, 1800 peptides CV<30%, n=3). It was noted that results from the 8 Da window setups were quite comparable (349 proteins; 1,794 peptides CV<30%, n=3) to that achieved using 12 Da windows (Figure 3F, 3G and 3H). As before, we investigated the impact of isolation window settings on the number of data point across the curve for the 11 spiked in iRT peptides and found the 8, 12, and 20 Da methods were associated with 12.3±3, 17.2±4.5 and 28.9±9.3 data points (n=3). Taken together, the 12 Da method was selected as optimal.^16^ We also evaluated our optimal methods for relative quantitative accuracy. Serial dilution curves were prepared, and samples were analyzed using the high-throughput method (concentration range: 31-500 ng) and mid-throughput method (concentration range: 31-2,500 ng). Relative peptide quantities were calculated by averaging the intensity response from all detected peptides across different concentration ranges (n=3). Figure 3I shows it was possible to establish a linear response across 31-500 ng for and 39-2500 ng for the high- and mid-throughput workflows in plasma, respectively. Using this analysis, an LLOD and LLOQ was determined for many of the peptides and proteins identified in the high- and mid-throughput analysis of plasma, depleted plasma and whole blood. See SuppFig 2 for summary and Supp Tables 1–14 for individual values.

### Precision of Standardized Workflow

Applying our optimized digestion protocol and DIA-MS methods, we compared the number of identifications along with the intra- and inter-day reliability of our workflow on three commonly analyzed biofluids: naive plasma, depleted plasma, and dried blood (See Table 1). For depleted plasma, the top 14 most abundant proteins were removed using the High Select Top 14 Abundant Protein Depletion Antibody Resin giving rise to a consistently produced subproteome that can be analyzed synergistically with naive plasma.^17^ Whole blood was collected using a Mitra device and dried prior to processing for LC-MS. To determine the analytical precision, 5 replicate samples of each biofluid were digested once a day for 3 consecutive days and analyzed using our high- and mid-throughput platforms. In the high-throughput method, cumulative frequency curves for the intra- and inter-day reproducibility showed the vast majority of peptides identified on individual days displayed CV<30% (93-96% plasma; 94-96% depleted plasma; 87-90% blood) (Figure 4A-C). Comparing across all 3 days, plasma has the greatest drop in peptide intensity reproducibility with only 74% of peptides achieving a CV<30% compared to 93% and 87% of peptides in depleted plasma and blood, respectively. In the mid-throughput platform, individual day peptide intensity reproducibility was high; greater than 85% of the peptides identified has CV<30% (89-93% plasma; 93-97% depleted plasma; 85-87% blood) (Figure 4D-E). Inter-day data, again, showed plasma to be the least reliable with only 67% of identified peptides below 30% CV compared to 90% for depleted plasma and 78% for blood.

**Table 1.** Summary of proteins and peptides identified in different biofluids using the standardized workflow.

|  |  | Protein IDs |  |  |  | Peptide IDs |  |  |  |
| --- | --- | --- | --- | --- | --- | --- | --- | --- | --- |
|  |  | Full | Reliable | Reproducible | Quantifiable | Full | Reliable | Reproducible | Quantifiable |
|  |  | Total Obs. | >2 Obs. / day | CV<30% | CV<30%+r<sup>2</sup>>0.8 | Total Obs. | >2 Obs. / day | CV<30% | CV<30%+r<sup>2</sup>>0.8 |
| **High-Throughput** |  |  |  |  |  |  |  |  |  |
| Blood | Multi-day | 365 | 236 | 189 | 121 | 823 | 530 | 466 | 320 |
|  | avg/day | 350±5 | 280±5 | 223±4 | 129±3 | 775±7 | 618±8 | 546±15 | 341±7 |
| Naïve Plasma | Multi-day | 240 | 161 | 133 | 98 | 1101 | 772 | 577 | 412 |
|  | avg/day | 215±3 | 181±5 | 163±4 | 117±4 | 1021±41 | 903±61 | 857±72 | 574±/18 |
| Depleted Plasma | Multi-day | 681 | 324 | 202 | 117 | 2190 | 1216 | 1108 | 321 |
|  | avg/day | 616±13 | 433±31 | 318±18 | 146±8 | 1996±53 | 1490±102 | 1376±92 | 353±31 |
| Combined Plasma | Multi-day | 754 | 378 | 266 | 144 | 2676 | 1547 | 1390 | 887 |
|  | avg/day | 674±15 | 486±29 | 383±15 | 169±6 | 2434±74 | 1875±127 | 1770±115 | 1003±42 |
| **Mid-Throughput** |  |  |  |  |  |  |  |  |  |
| Blood | Multi-day | 564 | 365 | 279 | 185 | 1572 | 1087 | 874 | 554 |
|  | avg/day | 518±5 | 426±15 | 353±14 | 209±7 | 1451±5 | 1238±27 | 1081±68 | 612±18 |
| Naïve Plasma | Multi-day | 572 | 256 | 168 | 152 | 2367 | 1416 | 1005 | 950 |
|  | avg/day | 515±45 | 381±82 | 303±59 | 249±38 | 2184±186 | 381±82 | 1691±327 | 1435±193 |
| Depleted Plasma | Multi-day | 462 | 321 | 281 | 198 | 2003 | 1618 | 1493 | 1051 |
|  | avg/day | 438±8 | 373±7 | 333±5 | 221±3 | 1992±25 | 1765±33 | 1678±38 | 1108±26 |

Total Obs. = total of the unique identifications observed either across all three days (black text) or the on individual days as avg/day (grey text; expressed as mean ± standard deviation). The other columns show counts meeting the stated threshold.

**Figure 4.**
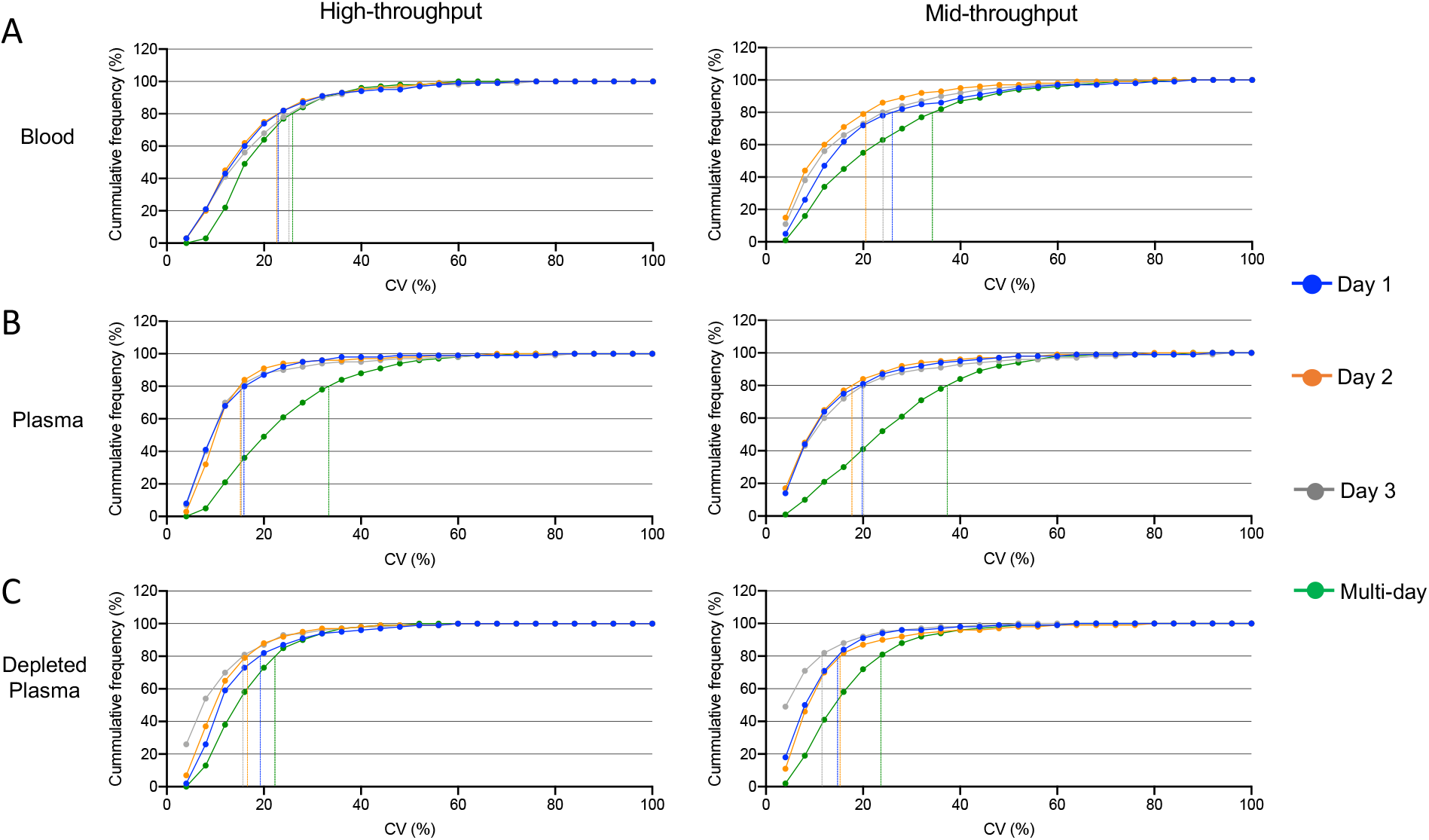
Evaluation of intra-day and inter-day precision in peptide quantification reproducibility. 5 replicates × 3 preparation days of robotic sample processing across three matrices: blood (**A**), naive plasma (**B**) and depleted plasma (**C**). Each set of sample preparation was acquired on two LC setups 25-min method (left panels) and 60-min method (right panels). Dashed lines are showing CV for 80% of peptides in each sample type and preparation day.

To compare the datasets produced from each biofluid and throughput method we established four levels of stringency (Table 1). The full proteome represents any proteins or peptides identified in any of the replicate day’s acquisitions. The reliable proteome are proteins observed in 3 or more runs on each of the 3 days. Reproducible identifications are those with a multi-day CV<30% and quantifiable are identifications meeting the reproducibility standard and having an r^2^>0.8 as determined by the linearity experiments. Comparison of the reliable proteomes found that depleted plasma has the largest number of proteins observed in the high-throughput method (324) while dried blood had the greatest number of proteins in the mid-throughput approach (365) (Figure 5A and C). It is notable that while most biofluids has an increase in reliable identifications with the longer separation time, depleted plasma was essentially constant. As an additional comparison, the high-throughput reliable identifications from naive and depleted plasma were combined; the non-redundant list was found to have a greater number of proteins than any of the individual mid-throughput analyses (See SuppTables 13-14).

**Figure 5:**
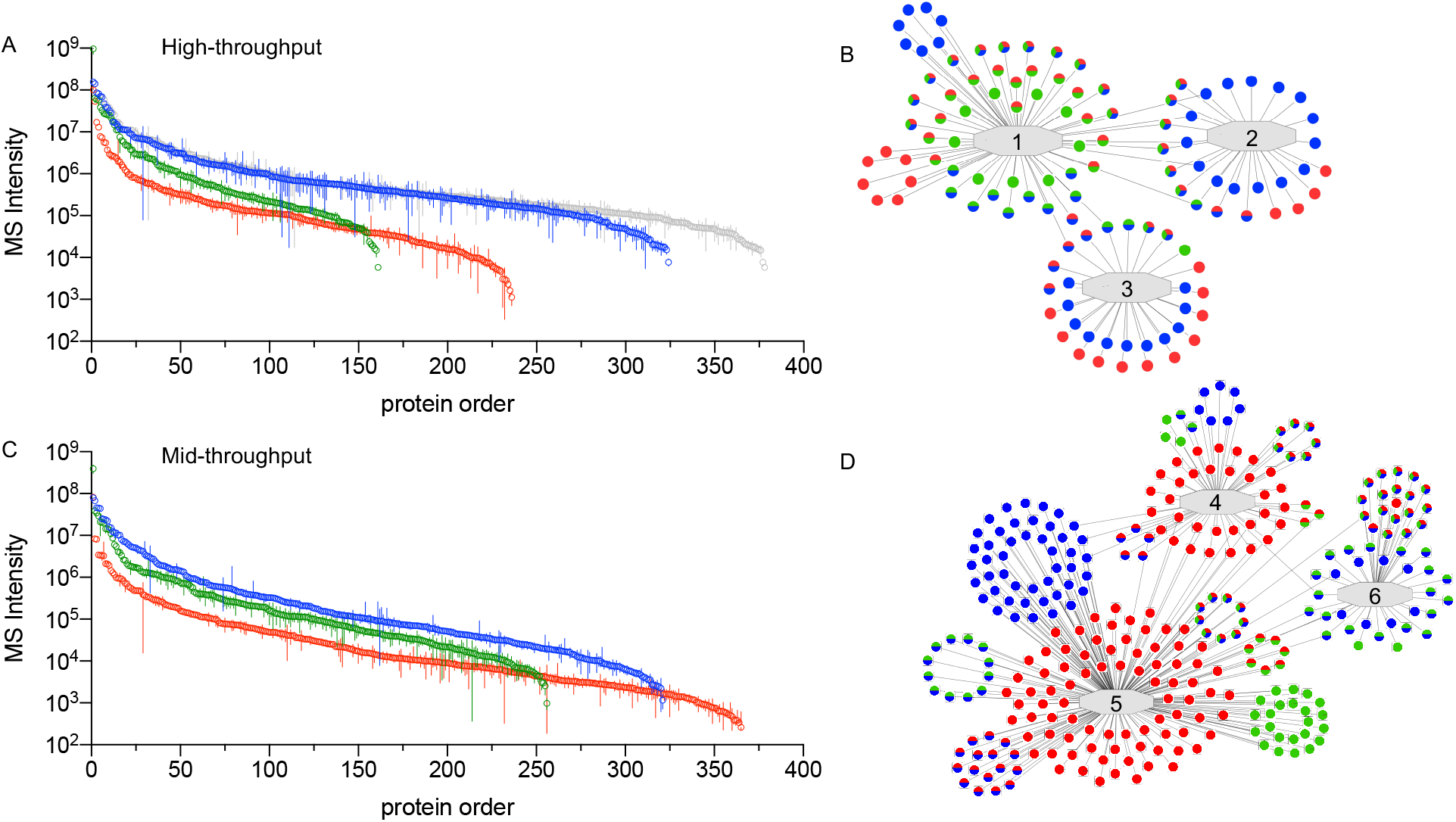
Comparisons of observed proteomes in standardized workflows. The number of reliable protein observations (>2/day) is shown for blood (red), naive plasma (green) and depleted plasma (blue) in the high-(**A**) and mid-throughput (**C**) workflows. Proteins were ordered based on the multi-day mean intensity (error bars=standard deviation). A non-redundant combination of the naive and depleted plasma is shown in grey. Examples of network analysis is shown for the high-(**B**) and mid-(**D**) throughputs. Networks: 1, adaptive immune response (GO:0002250); 2, ERK1 and ERK2 cascade (GO:0070371); 3, mitochondrial matrix (GO:0005759); 4, oxidoreductase activity (GO:0016491); 5, intracellular non-membrane bound organelles (GO:0043232); 6, complement and coagulation cascade (KEGG:04610). Dots indicate identified proteins (colors match panels **A** and **C**). Gene names were omitted for space. Fully annotated versions of panel (**B**) and (**D**) are presented in SuppFig 3 and 4 and for a complete listing of functional network assignments see SuppTables 15-16.

As a further characterization of the various proteomes, a pathway analysis was performed on the reliably identified proteome for each biofluid (Figure 5 B and D). In the high-throughput analysis proteins from each proteome were observed in many of the networks. Naive plasma has an enrichment in detected proteins involved in innate immunity while whole blood and depleted plasma had a greater enrichment in mitochondrial matrix proteins. The mid-throughput analysis of blood found enrichment of proteins involved in the oxidoreductase pathway while the complement and coagulation cascade were preferentially detected in the naive and depleted plasma analysis. See SuppTables 15 and 16 for a full listing of the functional pathways identified.

## Discussion

In this study we have developed and optimized a standardized workflow for three distinct blood biofluids, characterized the intra- and inter-day variability, and assessed the quantifiability of peptides and proteins detected. In keeping with advancements in MS-sample preparation workflows, our pipeline utilizes a robotic workstation for the core sample processing steps, namely: protein denaturation, cysteine reduction and alkylation, trypsin digestion, and peptide desalting. Maximizing the commonality in the workflow between biofluids simplifies the approach and increases the likelihood for reproducibility across samples, studies and institutions.

Historically, protein denaturation, enzymatic cleavage, and peptide desalting were considered to be especially vulnerable steps and are critical for effective sample preparation. A landmark study by *Borcehrs et al.* demonstrated that, while SDS predictably yields the most efficient digestion of the plasma proteome, TFE 50% (v/v), a volatile solvent, can be used in its place and provide comparable protein level data. Tryptic digestion of denatured proteins has been suggested to be maximal at elevated temperatures.^18–20^ Consistent with these reports, our data revealed incubation temperatures of 37 °C and 42 °C, as well as incubation times of 4 and 16 hr all supported reproducible proteolysis. Based on peptide numbers and amount of tryptic miscleavages, we selected parameters 4hr at 42 °C as the most time-effective protocol. There is a balance between plasma protein denaturation and deactivating the enzymatic activity of trypsin, TFE was diluted prior to addition of trypsin. Diluting TFE from 35 to 5% v/v instead of 10% substantially decreased the miscleavage rate by 13% and enhanced peptide recovery.

While an efficient digestion protocol helps overcome some of the hurdles associated with biomarker discovery in plasma, the vast dynamic range and missing data especially for low abundant proteins still poses challenges. The use of peptide libraries with DIA-MS reduces missing data but generates very complex chimeric spectra posing a challenge to downstream data analysis. Narrower isolation windows with DIA-MS can partially reduce this complexity provided that they are not taxing on the cycle time required to cover the typical mass range (e.g. 350-1400 m/z).^21^ Chromatography methods using wider diameter columns (300 µm) and faster flow rates are replacing traditional nanoflow techniques despite the trade off with sensitivity.^16^ The recently developed LC system, Evosep One, provides disposable trap columns (Evotips) and rapid low flow gradients (3-88 min), and this system is designed to drive throughput towards an analysis of 15-300 samples per day.^22^ We took advantage of the 60 sample-per-day workflow (21 min gradient) offered by Evosep to support our high-throughput set up. For our mid-throughput platform, we adhered to a more conventional chromatographic set up using LC column of 3 µm particle size, polar C18 with pore size 100 Å and 150 x 0.3 mm dimensions (Phenomenex, Torrance, CA). By screening DIA m/z windows on each system we found optimal settings that maximized reproducibility. The shorter gradient performed better with slightly larger mass windows (21 Da) compared to the longer gradient (12 Da) which is consistent with the relative complexity of the sample at those flow rates. The result is 2 robust systems for practical applications: high-throughput plasma biomarker discovery and mid-throughput deep profiling analysis.

Precise quantification of low-abundance proteins in plasma remains a challenge, which can be addressed using depletion strategies that simplify the plasma sample proteome complexity. These procedures are themselves not devoid of limitations, chief among them being the additional sample handling steps required that can compromise analytical reproducibility. Furthermore, traditional depletion protocols are performed on a per-sample basis using liquid chromatography systems, and this timely process requires large volumes of starting material.^17,23,24^ Of note, our described 96-well plate protocol utilizes antibody slurry and starts with a plasma volume of only 10 μl which is consistent with the small sample volumes available in many clinical and translational studies.

Collection of dried blood by a self-administered minimal invasive finger prick device removes several barriers associated with the delivery of remote medical support.^25,26^ We have amended our original workflow for processing the 10 μl of blood absorbed and dried onto a Mitra^®^ microsampling device to conform with the two new plasma protocols, showing that dried blood, while different in composition to naive and depleted plasma, can be analyzed with similar levels of reproducibility and quantifiability compared to the other two more prominent sample types in biomarker discovery studies. It was noted that our high-throughput assay supported greatest IDs from depleted plasma while our mid-throughput assay yielded the largest number of proteins from whole blood. Depleted plasma represents the cleanest proteome in terms of sample complexity while whole blood is subjected to the fewest pre-processing steps. Our loading experiments revealed the high-throughput assay was associated with reduced loading capacity compared to the mid-throughput assay. We therefore hypothesis that background matrix resulting from hemolysis of red blood cells had a greater impact on the high-throughput assay in comparison to the mid-throughput assay. In contrast, since it is associated with the fewest pre-processing steps it was not surprising to find whole blood yielded the most reliable proteome when analyzed by the mid-throughput assay. To fully address this interference studies will be carried out as a part of future work.

The results from our precision study demonstrated that, with only modest adaptations, our plasma workflow can be used to reproducibly prepare samples derived from various clinical matrices (e.g. depleted plasma and whole blood). Table 1 summarizes the performance of the different biofluids indicating all can provide reliable and reproducible analysis. Of note, we observed that by combining the naive and depleted plasma analyses from the high-throughput workflow produced a comparable number of reliable identifications compared to an individual mid-throughput analysis (Figure 5A). This is significant as two 25-min runs on the Evosep One can be completed in 50 mins while a 60-min run on the Ultimate 3000 system requires 70 mins to complete, including a 12 min blank run. Using the combined plasma approach, depending on the needs of the experiment, it could be possible to achieve a similar depth of analysis in approximately 70% the instrument time.

In summary, our streamlined process is associated with minimal hands-on time, fast turnaround, and excellent intra- and inter-day stability. The high-throughput platform supported reproducible detection of more than 74% of peptides in all matrices assed. While the mid-throughput platform supported reproducible detection of greater than 67% of peptides across all matrices. Of special interest, we observed that depleted plasma’s inter-day precision was the most stable, with more than 90% of peptides reproducibly detected across days. We therefore anticipate that our workflow can be easily transferred to other research institutions to help support cost-effective broad scale multi-center biomarker studies. Our broad vision for this platform is that it will help propel putative markers into clinical use.

## Acknowledgements

The authors would like to thank all of the staff at the CSMC biobank and precision biomarker labs for their help collecting and aliquoting samples.

Sources of funding

This work is supported by the Erika Glazer Covid 19 Fund to J.V.E. and S.C. and NIH grant U01 NS115658-01 to J.V.E.

