## Supplementary material for "Standardized workflow for precise mid- and high-throughput proteomics of blood biofluids": Online Supplement

### Supporting Information

### Methods

#### Loading of Mitra tips

Pool of human whole blood (with $K_2EDTA$ preservative) was obtained from Bioreclamation IVT (Chestertown, MD, US). All blood pools were stored at 4°C. Mitra® microsampling devices (Neoteryx, Torrance, CA, US) with 10 µL volumes were loaded with blood by dipping the tips into aliquoted 10uL-volume droplets of whole blood. Care was taken that the Mitra tips only touched the liquid surface and held at the droplet surface until fully colored red ensuring a complete fill. The filled Mitra tips were allowed to dry for at least 4 h at room temperature and stored until further processing in a closed container in the presence of a desiccant at -80°C.

#### Automated Desalting

Digested samples (300 ul) were mixed with 850 ul of 2% phosphoric acid, 0.1% formic acid. The desalting plate was activated with addition of 1 ml of 100 % methanol and equilibrated 3 times with the addition of 1 ml of 0.1% formic acid. 1150 ul of the acidified sample was applied to the plate and then washed with 3 applications of 0.1 % formic acid. Peptide desalting was performed using Oasis 30 µm HLB 96-well Plate (Waters Co.) was adapted for the Beckman i7 automated workstation.[1] The positive pressure apparatus (Amplius Positive Pressure ALP, Beckman Coulter) was mounted on the left side of the i7 workstation deck and the solid phase extraction pipetting steps (desalting) were carried out by i7 pipetting heads and controlled by Biomek Software (v5.1). For MeOH activation, filter pressure was set at 100 mBar and increased to 250 mBar over 1 min and 30 seconds. For washing and elution steps, the pressure started at 100 mBar and increased 50 mBar every 15 seconds to a final filter pressure of 550 mBar over 2 mins and 30 seconds. For each protocol a clamp pressure of 3000 mBar at the beginning and end. The sample was eluted with 0.5 ml of 50 % acetonitrile, 0.1% formic acid and evaporated to dryness and stored at -80ºC. At the time of analysis, peptides were resuspended in a 0.1% formic acid solution.

#### DDA LC-MS/MS

For DDA experiments, an Evosep 30 sample per day workflow which uses a 15 cm x 3um C18 column (Evosep) and a 45 min gradient was implemented. Full MS resolution was set to 60,000, scan range was 350-1,250 m/z 200 AGC target was 300%. For MS2, resolution was 15,000, isolation width was 1.2 m/z, HCD energy was 28% and AGC target was set to 75 %.

#### High- and Mid-throughput DIA LC-MS/MS

DIA analysis was performed on an Orbitrap Exploris 480 (Thermo) instrument. For the high throughput workflow, the instrument was interfaced with an easy nano spray ion source coupled to an Evosep One. Peptides were separated on a preformed gradient (ranging from 5 - 35% organic phase) on a C18 column (8 cm, 3 µm) over the course of 21 mins at a flow rate of 1000 nl/min. Source parameters included spray voltage at 2000 kV, capillary temp of 275 °C and RF funnel level of 40. MS1 resolutions were set to 120,000 and AGC was set to 300% with ion transmission of 45 ms. Mass range of 350-1400 and AGC target value for fragment spectra of 300% were used. Peptide ions were fragmented at a normalized collision energy of 28%. Fragmented ions were detected across 50 DIA windows of 21 Da with an overlap of 1 Da (Adapted from [2]). MS 2 resolutions was set to 15,000 with an ion transmission time of 22 ms. All data was acquired in profile mode using positive polarity.

Mid-throughput DIA analysis was performed on an Orbitrap Exploris 480 (Thermo) instrument interfaced with a flex source coupled to an Ultimate 3000 ultra high-pressure chromatography system with 0.1% formic acid in water as mobile phase A and 0.1% formic acid in acetonitrile as mobile phase B. Peptides were separated on a linear gradient of 1-27% B organic phase for 45 min, 27-44% B for 15min on a C18 column (15 cm, 3 µm) over the course of total 60 mins at a flow rate of 9.5 ul/min. Between every sample the column was washed with a 10 min blank where the organic phase was increased to 98% and then re-equilibrated at 1% B for 2 mins. Source parameters included; spray at 3000 kV, capillary temp of 300 °C and an RF funnel level of 40. MS1 resolutions was set to 60,000 and AGC was set to “standard” with ion transmission of 100 ms. Mass range of 400-1000 and AGC target value for fragment spectra of 300% was used. Peptide ions were fragmented at a normalized collision energy of 30%. Fragmented ions were detected across 50 DIA non-overlapping 12 Da precursor windows. MS2 Resolution was set to 15,000 with an ion transmission time of 25 ms. All data is acquired in profile mode using positive polarity.

MS data and analysis has been deposited in the Panorama public repository.

#### Bioinformatic Data Analysis

LC-MS/MS data were visually inspected using XCalibur software (4.3.73.11). DIA MS raw files were converted to mzML, the raw intensity data for peptide fragments were extracted from DIA files using the OpenSWATH workflow and searched against the Human Twin population plasma peptide assay library. Retention time predictions were made using the data from Biognosys iRT Standards spiked into each sample. Target and decoy peptides were then extracted, scored and analyzed using the mProphet algorithm to determine scoring cut-offs consistent with 1% FDR. Peak group extraction data from each DIA file was combined using the 'feature alignment' script, which performs data alignment and modeling analysis across an experimental dataset. The total ion current (TIC) associated with the MS2 signal across the chromatogram was calculated for normalization using in-house software. This 'MS2 Signal' for each file was used to adjust the transition intensity of each peptide in a corresponding file. Normalized transition-level data was then processed using the mapDIA software to perform quantitation at the peptide and protein level.

A full description of the data analysis and all peptide and proteins identifications including reproducibility and linearity characterizations can be found in the supporting information and Supplemental Tables 1-14. The coefficients of variation (CV) for protein or peptide intensities were determined if there were at least 3 out of 5 observations on each day. This threshold was required for all 3 days to determine a multi-day CV. Observed lower limit of detection (LLOD) for a protein or peptide based on the linearity experiment and determined by the lowest concentration of sample where the protein or peptide is detected in all 3 replicates with a CV<20%. Observed lower limit of quantitation (LLOQ) for a protein or peptide, was determined by the lowest concentration of sample where the protein or peptide is detected in all three replicates with an $r^2>0.8$, CV<20% and a target deviation>0.2. Lower limit of detection and quantifiability were also estimated by LLOD/Q = SD residuals/slope*(3.3 or 10) for proteins that displayed a $r^2>0.8$ in a linear regression across at least 3 consecutive loading points concentrations.[3]

Functional pathway characterization and visualization was performed using ClueGO Ontology Analysis via PINE (Protein Interaction Network Extractor).[4,5] Visualization of significantly enriched terms in whole blood compared to native and depleted plasma in high- and mid-throughput methods. PINE consolidates protein interactions information from STRING and GeneMANIA to create a single, unified network.[6,7] Category type analysis was performed in PINE to plot protein distribution across 3 categories: whole blood, native and depleted plasma from the high- and mid-throughput workflows.


### Supplementary Tables

Supplementary Tables 1-4: Characterization of proteins and peptides detected in dried blood using a high- and mid-throughput workflows.

Supplementary Tables 5-8: Characterization of proteins and peptides detected in naïve plasma using a high- and mid-throughput workflows.

Supplementary Tables 9-12: Characterization of proteins and peptides detected in depleted plasma using a high- and mid-throughput workflows.

Supplementary Tables 13-14: Characterization of non-redundant proteins and peptides detected in a combination of the naïve and depleted plasma analyses from the high-throughput workflow.

Supplementary Tables 15-16: Functional network assignments for proteins reliability detected in blood, plasma and depleted plasma in the high- and mid-throughput workflows.

### Supplementary Figures

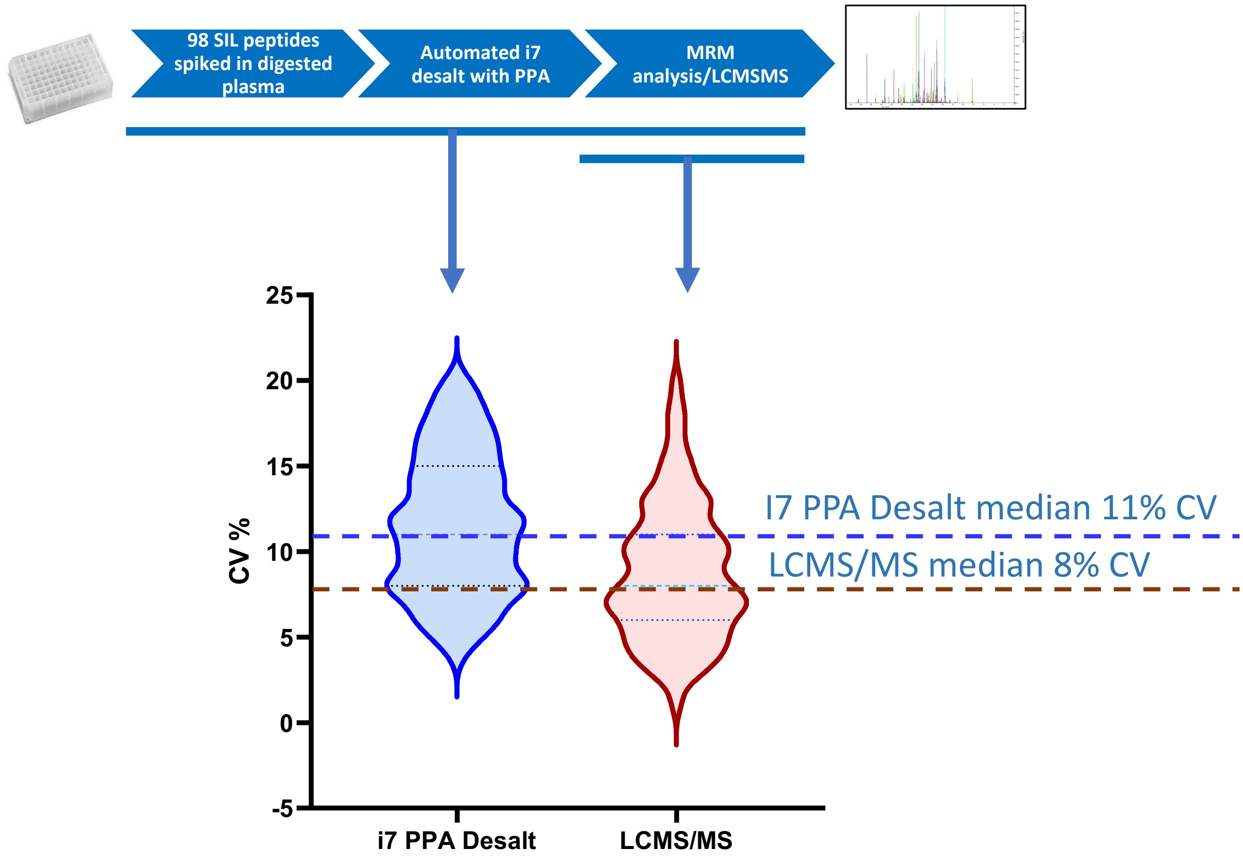


**Supp Figure 1: The Reproducibility of automated i7 desalt (solid phase extraction) method.** The precision of total desalt workflow is comprised of both %CV from i7 workstation desalting and subsequent targeted LC MS/MS. Stable isotope labeled internal standard (SIL) peptides spiked in digested human plasma were used to evaluate reproducibility of the method. The precision was determined from 18 wells/samples desalted with a HLB plate, processing 98 SIL internal standards in trypsin digested plasma in representative solid phase extraction experiment and %CVs for LC MS/MS analysis and automated desalt processing were accessed with a highly multiplexed MRM assay (J Proteome Res 2018. doi: 10.1021) (PMID: 32391810). Briefly, tryptic plasma peptides and 98 internal standards were aliquoted into 18 wells and desalted by the i7 workstation. The desalted 98 SIL heavy peptides quantified by the MRM analysis with a Prominence UFLCXR HPLC system (Shimadzu, Japan) with a Waters Xbridge Peptide column coupled to a QTRAP® 6500 with a Turbo V source. Analyst® software (version 1.6.2 for the QTRAP 6500 was used to control the LC–MS system and for data acquisition. All MRM data were processed using MultiQuant™ 2.1 Software (SCIEX). The automated i7 desalt method demonstrated good reproducibility. The total %CV ranged from 4%-20% (with median 11% CV) for 247 transitions (representing 98 peptides and 53 proteins) after i7 desalting of SILs using targeted MRM analysis. The baseline of LC-MS errors in the MRM analysis were calculated from 7 repeated LC-MS injections. The identical 247 transitions showed %CV ranged from 1%-20% (with median 8%) for LCMS/MS.

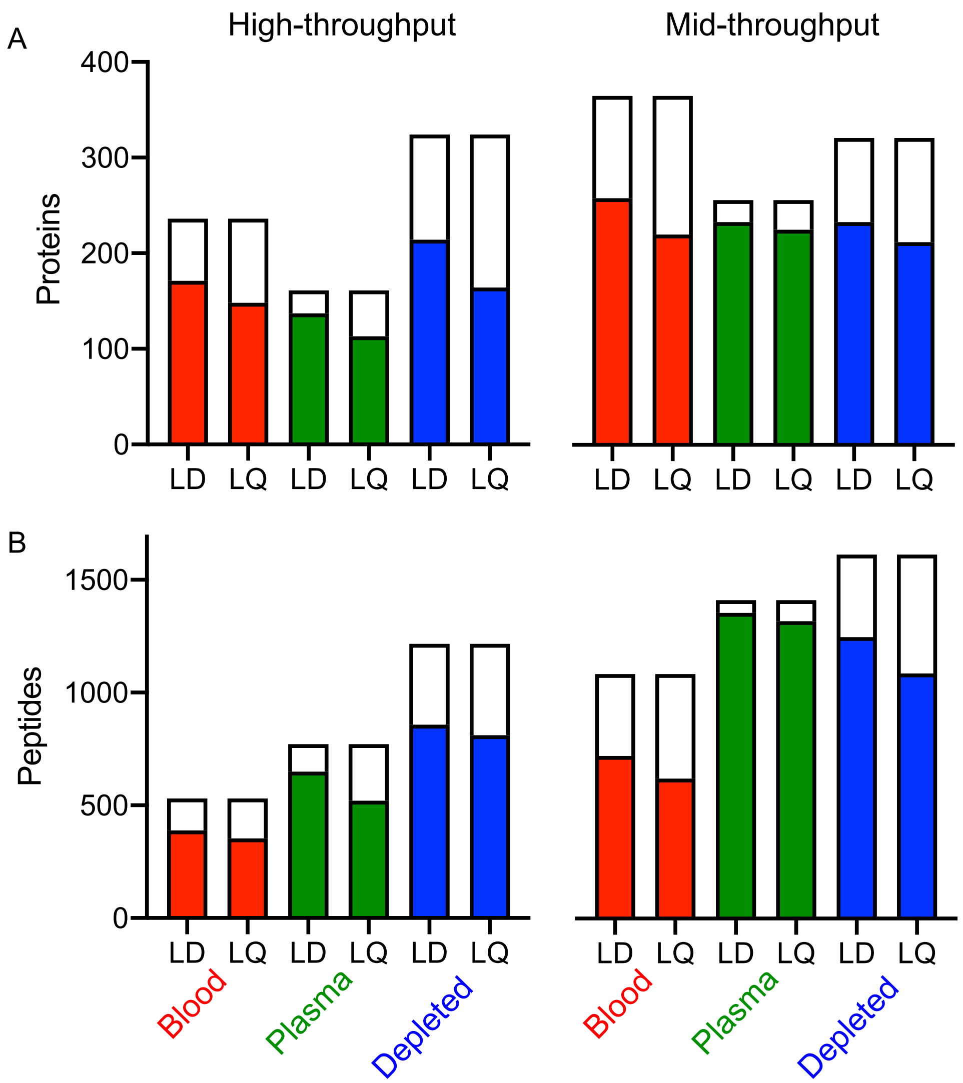


**SuppFigure 2. Determination of LLOD (LD) and LLOQ (LQ) for the reliable proteome in each biofluid using the standardized workflows.** LLOD and LLOQ were determined from linearity experiments for proteins (A) and peptides (B) in the high-(left) and mid-(right) throughput workflows. Total bar height indicates number of proteins or peptides with at least 3 observations per day. The filled bars indicate the number of proteins or peptides where an LLOD or LLOQ was determined. See Methods for description of LLOD/Q determination.

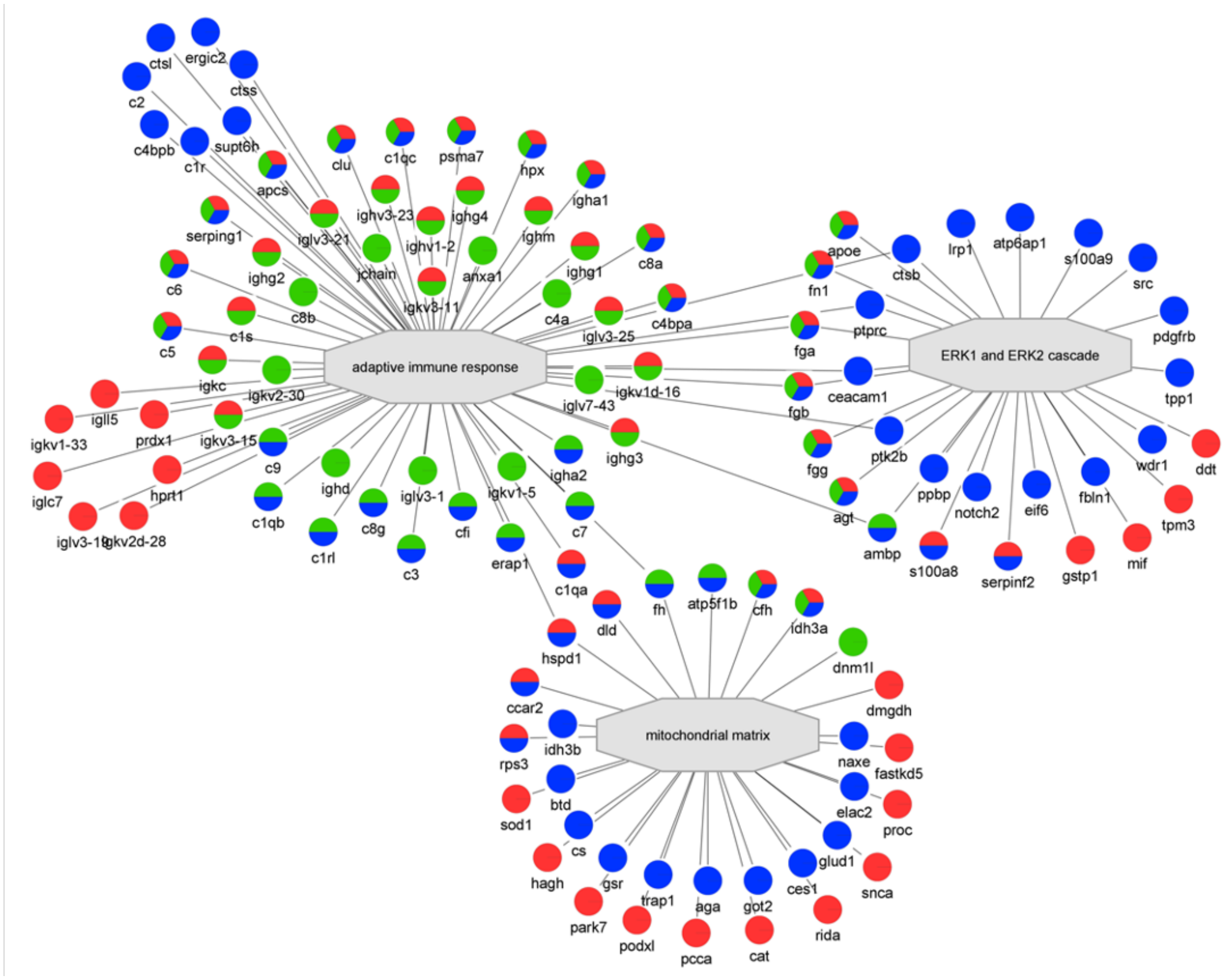


**SuppFigure 3. Network analysis for high-throughput workflow.** An example of the PINE category type analysis is shown for reliably identified proteins (at least 3 observations on each of the 3 days) from the whole blood (red), naive plasma (green) and depleted plasma (blue). The central grey nodes denote term of enrichment. See SuppTables 15 for a complete listing of functional network assignments.

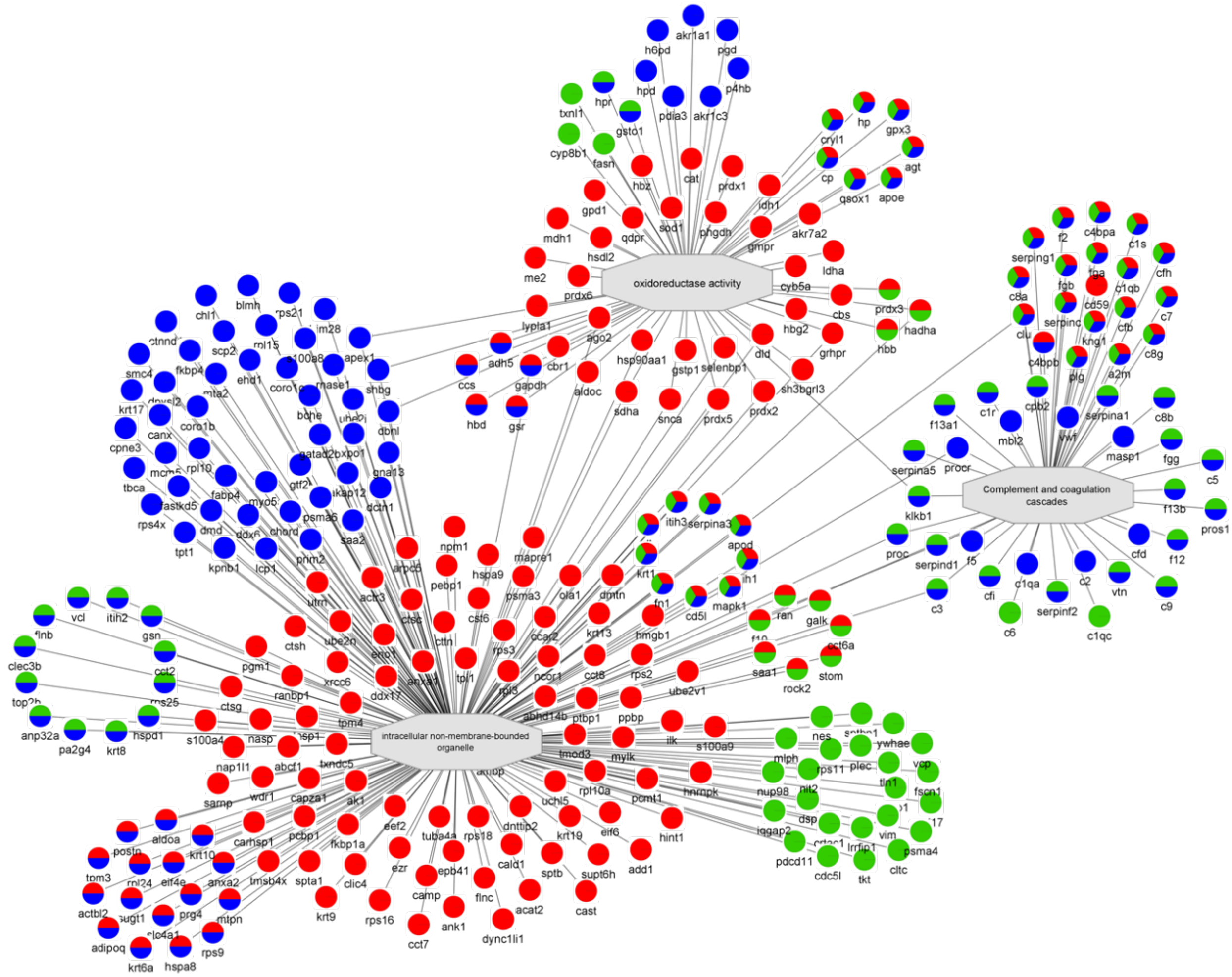


**SuppFigure 4. Network analysis for mid-throughput workflow.** An example of the PINE category type analysis is shown for reliably identified proteins (at least 3 observations on each of the 3 days) from the whole blood (red), naive plasma (green) and depleted plasma (blue). The central grey nodes denote term of enrichment. See SuppTables 16 for a complete listing of functional network assignments.
